# Quantifying topography-guided actin dynamics across scales using optical flow

**DOI:** 10.1101/753681

**Authors:** Rachel M. Lee, Leonard Campanello, Matt J. Hourwitz, Ava Omidvar, Phillip Alvarez, John T. Fourkas, Wolfgang Losert

## Abstract

Periodic surface topographies with feature sizes comparable to those of *in vivo* collagen fibers are used to measure and compare actin dynamics for two representative cell types that have markedly different migratory modes and physiological purposes: slowly migrating epithelial MCF10A cells and polarizing, fast migrating, neutrophil-like HL60 cells. Both cell types exhibit reproducible guidance of actin waves (esotaxis) on these topographies, enabling quantitative comparisons of actin dynamics. We adapt a computer-vision algorithm, optical flow, to measure the directions of actin waves at the submicron scale. Clustering the optical flow into regions that move in similar directions enables micron-scale measurements of actin-wave speed and direction. Although the speed and morphology of actin waves differ between MCF10A and HL60 cells, the underlying actin guidance by nanotopography is similar in both cell types at the micron and sub-micron scales.

---

Understanding the rearrangements of the cytoskeleton is essential to developing a complete picture of dynamic cellular processes such as migration, division, and differentiation. Cytoskeletal dynamics, and in particular actin dynamics, have been shown to be important for the growth of cell junctions and focal adhesions^1^ and for immune-cell activation^2^. The formation of actin waves through directional polymerization and depolymerization of filaments drives many types of cell migration^3^, and has been associated with the establishment of polarity in a variety of cell types^4^.

An important modulator of actin dynamics is the extracellular environment. Physical and chemical characteristics of the extracellular environment, such as rigidity, biochemical composition, and topography, have been shown to influence actin dynamics and associated cell behavior^5–8^. One mechanism for this modulation is mechanosensing via focal adhesions^9^. In addition, actin waves respond when cells encounter obstacles^10^. It has also been established that ridges of width comparable to fibers in the extracellular matrix (ECM) can alter actin dynamics significantly^9,11,12^ and bias the localization of focal adhesions^13,14^. Thus, *in vivo*, the topography of the ECM, such as collagen networks^15,16^, is likely to modulate actin dynamics.

Periodic nanotopographic surfaces provide the opportunity to obtain systematic data on the modulation of such intracellular dynamics. In prior work we have shown that actin waves can be nucleated near, and guided along, periodic nanotopography, in a phenomenon termed esotaxis. Actin-wave guidance has been observed in cell types that exhibit distinct physiological functions and migration phenotypes, including *Dictyostelium discoideum*^11,12^, neutrophil-like HL60 cells^12^, B cells^9^, and breast-cancer cell lines^17^. However, there are clear differences in the responses of each of these cell types to nanotopography. For example, although both *D. discoideum* and HL60 cells exhibit esotaxis, these two types of cells have been found to move preferentially in different directions on specific nanoscale asymmetric sawtooth textures^12^. Furthermore, different breast-cancer cell lines preferentially move in different directions on asymmetric sawtooth nanotopography^17^.

Here we introduce a method for performing quantitative measurements of the influence of nanotopography on intracellular dynamics at both the submicron and micron scales. This approach enables the detection of subtle differences in cytoskeletal dynamics and allows for in-depth analysis of both the differences and similarities of these dynamics across cell types and phyla. Our method of quantification of actin dynamics across scales is based on optical flow, an image-analysis technique developed in the fields of robotics and navigation control that uses changes in pixel intensities to detect motion in image sequences^18,19^. Here we use this technique on fluorescence micrograph time series to measure the spatiotemporal dynamics of actin with submicron precision. Clustering of the optical-flow data further allows us to quantify actin dynamics on the micron scale. Thus, this optical-flow-based analysis enables the identification of similarities and differences between esotaxis in neutrophil-like HL60 cells and human breast epithelial MCF10A cells across length scales.

## RESULTS

Esotaxis has been observed in a wide range of cell types that are known to respond to their *in vivo* microenvironment through processes such as directed migration or immune-system activation^9,11,12^. More recently, esotaxis has also been observed in epithelial cells, which are less motile^17^. Here we contrast the actin dynamics of epithelial MCF10A cells with those of neutrophil-like HL60 cells.

LifeAct-GFP-labeled epithelial MCF10A cells were plated on a 900 µm × 900 µm region patterned with parallel nanoridges with a spacing of 1.5 µm, as well as on the surrounding flat region. Confocal imaging near the surface revealed distinct actin morphologies on nanoridges as compared to the flat region (Figure 1). On the flat region the phenotype is a common one for these cells on such surfaces, with a broad lamellipodium at the cell front and stress fibers throughout the cell body (Figure 1a). In contrast, MCF10A cells on nanoridges exhibit actin streaks aligned with the ridges throughout the cell area (Figure 1b). The local nature of the response of actin to surface texture is illustrated in Figure 1c, which shows a cell that is partially on the nanoridges and partially on the flat region. On the nanoridged region, the cell shows the same actin streaks as a cell that lies fully on a ridged surface, whereas the same cell maintains a broad lamellipodium on the flat region.

**Figure 1:**
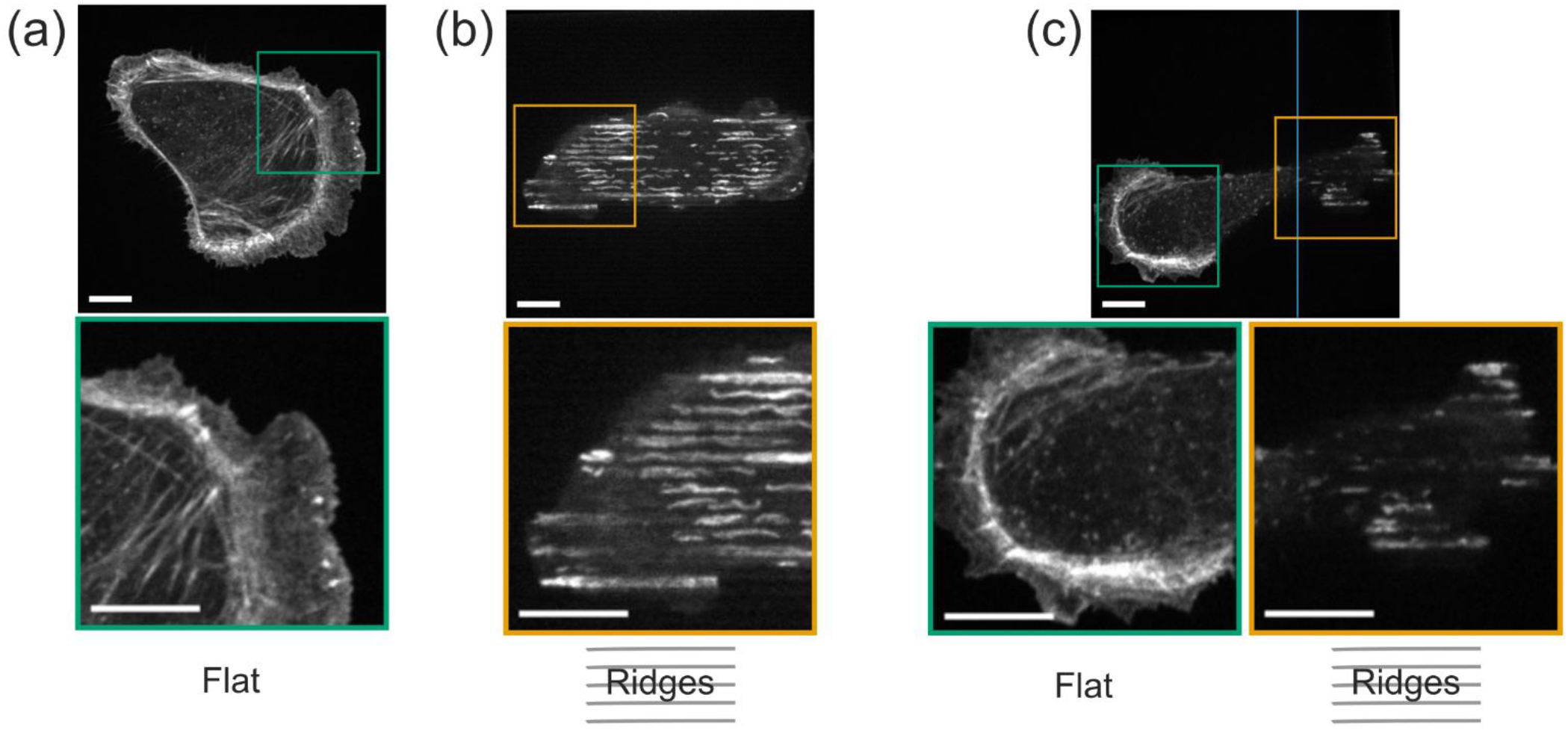
Surface topography prompts distinct actin morphology. The actin cytoskeleton of an epithelial MCF10A cell on (a) a flat surface has an actin morphology that is distinct from (b) that of a cell on a nanoridged surface. (c) A cell that is partially on a nanoridged region and partially on a flat region exhibits local actin morphologies that are driven by the underlying topography. The blue line in (c) indicates the boundary between the flat region and the nanoridged region. All scale bars are 10 µm. See also **Supplemental Movie 1**.

Ridged and flat regions also engender distinct actin dynamics. Kymographs can be used to visualize dynamics in a region of interest in one spatial direction over time. The left side of Figure 2 compares an MCF10A cell on a flat region with one on a nanoridged region. The cell on the nanoridged region shows actin streaks that are characteristic of esotaxis. Actin kymographs from two perpendicular regions (Figure 2a) in an MCF10A cell on a flat surface show oscillatory dynamics in all directions at the cell boundary. These oscillations in the kymographs indicate the presence of fan-like protrusions and retractions across each region over 30 min (Figure 2b). In contrast, on the nanoridged region, the actin dynamics parallel and perpendicular to the ridges are different (Figure 2c). Parallel to the nanoridges, MCF10A cells show oscillatory actin dynamics. As shown in the bottom left of Figure 2c, a representative kymograph of a region perpendicular to the ridges shows actin structures that persist for tens of minutes and do not move perpendicular to the ridges. This behavior is typical for kymographs perpendicular to the ridges, although perpendicular motion is observed in some cases as discussed below.

**Figure 2:**
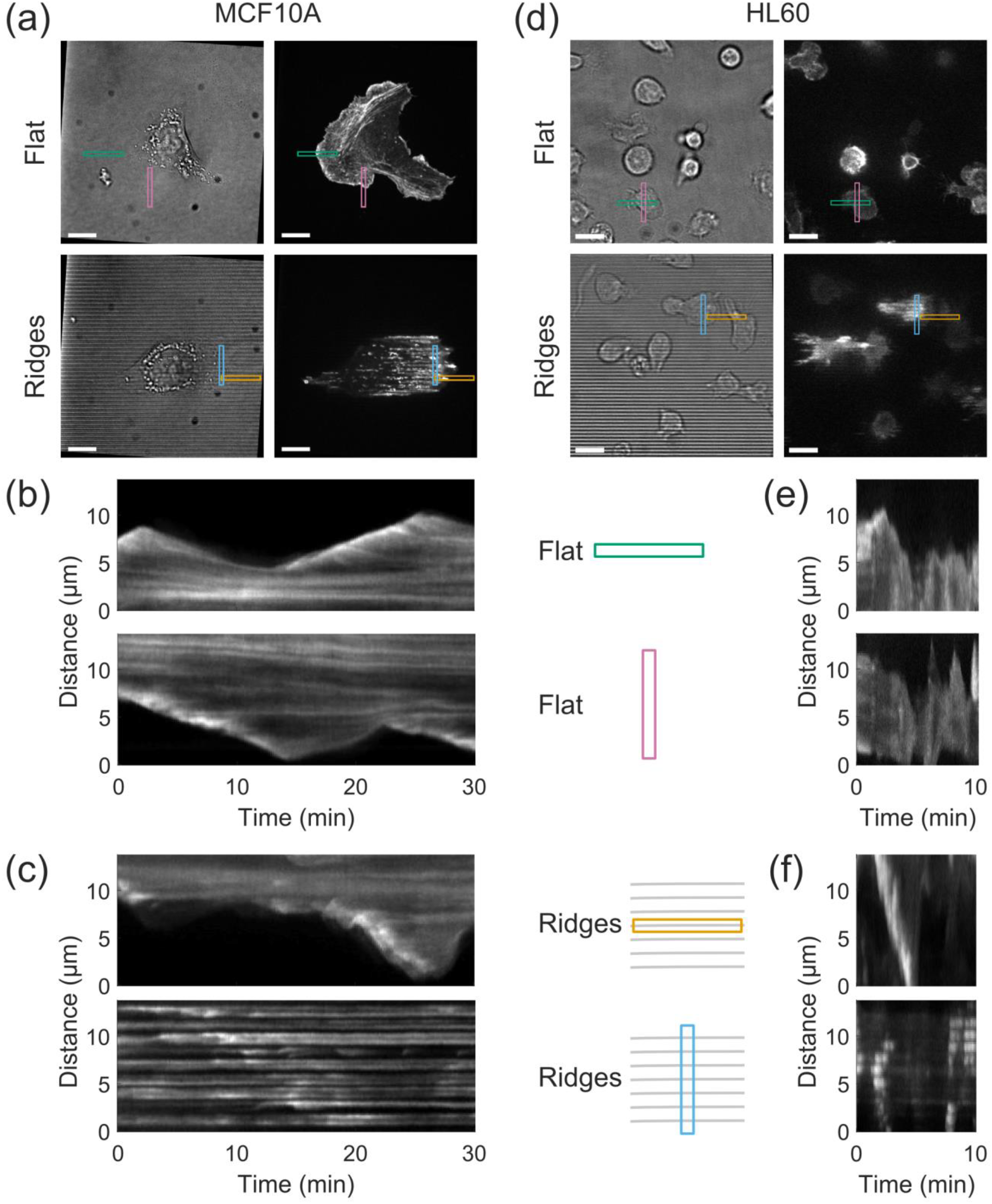
Surface topography leads to distinct actin morphodynamics. (a) Optical micrographs of MCF10A cells in (left) bright-field and (right) fluorescence on (top) a flat region and (bottom) a nanoridged region. All scale bars are 10 µm. Kymographs for the areas denoted in (a) are shown in (b) for the flat region and in (c) for the nanoridges. (d), (e), and (f) are the same as (a), (b), and (c), respectively, but for HL60 cells. The dynamics in (a, d) are shown in **Supplemental Movies 2 and 3**, respectively.

The behavior of motile, neutrophil-like HL60 cells on flat and nanoridged regions is illustrated in Figure 2d-f. Figure 2d illustrates the regions from which kymographs were generated. In HL60 cells on flat regions actin is concentrated near the cell front. This localization is largely preserved on the ridged surfaces, although the morphology of the actin changes such that streaks of actin are aligned with the ridges. On flat surfaces, the HL60 cells show regions of protrusions and retractions (Figure 2e) similar to the actin dynamics seen in MCF10A cells (Figure 2b). We note that protrusions occur on the scale of seconds in HL60 cells and on a scale of minutes in MCF10A cells. Kymographs of the HL60 cells in the direction parallel to the nanoridges show protrusive dynamics, although often in the form of a single persistent wave (Figure 2f, top), in contrast to the oscillatory behavior seen on flat surfaces (Figure 2e). A representative kymograph of an HL60 cell in the direction perpendicular to the nanoridges shows streaks (Figure 2f, bottom) that indicate that actin waves do not move perpendicular to the ridges, but the streaks are shorter in duration than those in a typical MCF10A cell (Figure 2c). This behavior is typical for kymographs of actin in HL60 perpendicular to the ridges. Unlike in MCF10A cells, in which actin streaks on the ridges localize throughout the cell (Figure 2c, bottom), in the HL60 cells the streaks occur near the cell front (Figure 2f, bottom). Groups of actin streaks propagate together at the front of the HL60 cells, suggesting that there may be large-scale organization of actin dynamics (spanning many ridges) in these cells. It is unclear whether there is large-scale organization of actin dynamics in the MCF10A cells.

As shown in **Supplemental Movies 2 and 3**, the full range of actin dynamics is more complex than is revealed by kymographs. MCF10A cells on the ridged regions exhibit actin dynamics throughout the substrate contact area, whereas actin dynamics on flat surfaces are largely confined to the cell boundary. These movies show that, in both cell types, nanoridges stimulate reproducible, dynamic, linear actin structures.

Time-lapse fluorescence images of actin waves are difficult to interpret by visual inspection or kymographs alone, because the observed dynamics arise from a complex spatio-temporal concentration field. To measure these wave-like dynamics quantitatively, we must first define a wave (size and shape) and then capture its propagation (splitting, recombination, and changes in direction). Here, we address these challenges by introducing an automated approach to quantify actin-wave dynamics across length scales for unbiased comparison in different cell types and extracellular environments.

Our method is based on a computer-vision algorithm from robotics and navigation control called optical flow^18,19^, which provides pixel-based information about the direction and magnitude of intensity flux in a series of time-lapse images. Fields of optical-flow vectors are calculated by integrating changes of intensity in space and time, as shown schematically in Figure 3. For example, two images of a migrating HL60 cell taken 8 s apart are shown with changes in time highlighted by a green-to-magenta montage (Figure 3a). The magenta region indicates growth of the actin front (which, as expected, occurs at the leading edge of the cell), and the green region indicates a decrease in actin intensity.

**Figure 3:**
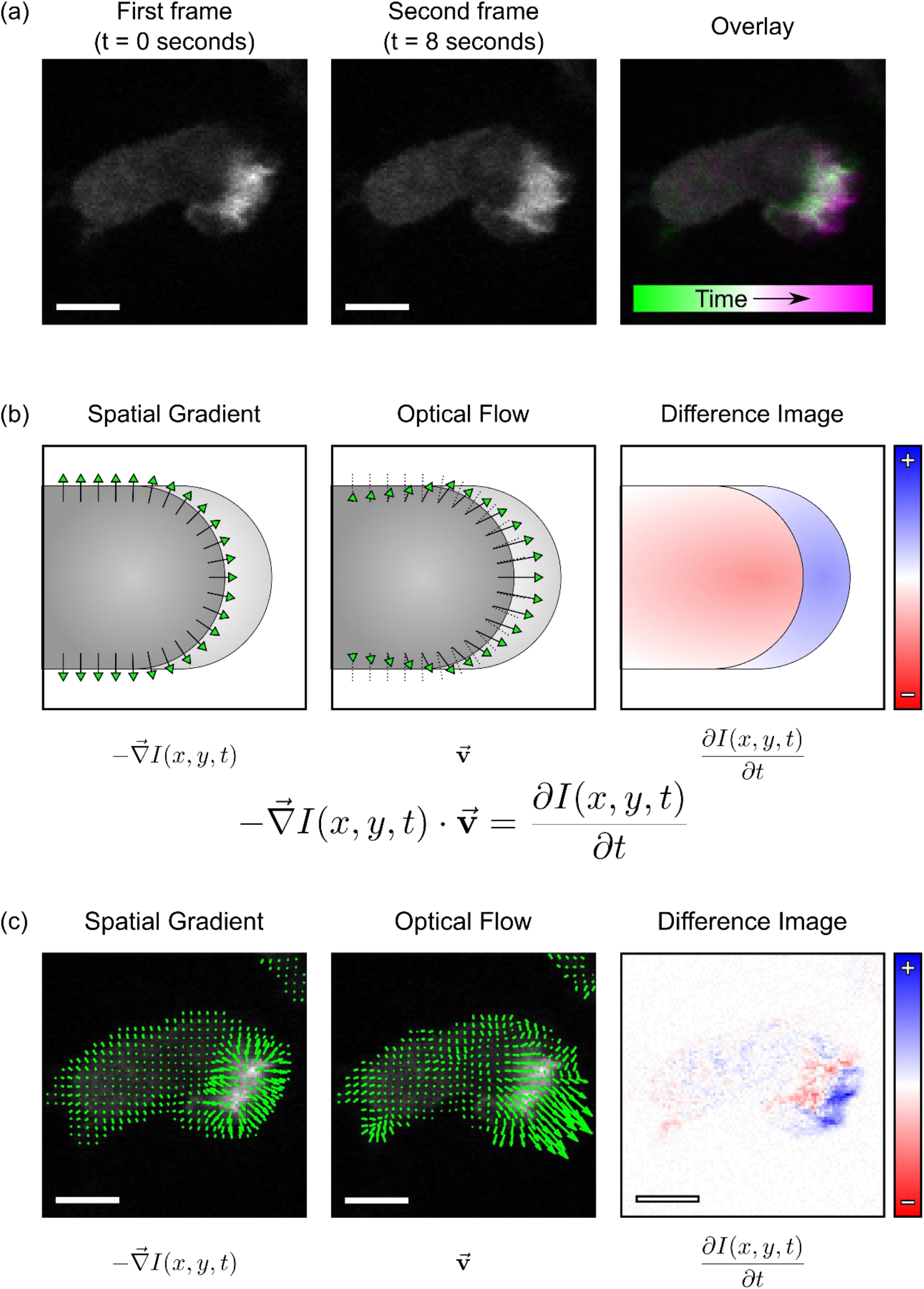
Optical-flow calculations capture the dynamics in movies of actin fluorescence. (a) Two frames of a representative HL60 cell obtained 8 s apart and a merged image show the dynamics of the cell’s behavior over time. The schematic in (b) illustrates how the procedure used to carry out optical-flow calculations combines the spatial gradient of an image (left) and the difference image/temporal gradient (right) to yield the optical-flow vector field (center). These calculations are applied to the images in (a) and shown in the images of (c). The spatial gradient field (left) and temporal gradient (right) result in the output optical-flow vector field (center). Blue pixels in the right panel of (c) indicate a positive change (increase) in the pixel brightness from the first frame to the second frame, and red pixels in the right panel indicate a negative change (decrease) in the pixel brightness from the first frame to the second frame. All scale bars are 5 µm.

The general objective of calculating optical flow is to solve for the unknowns Δ*x* and Δ*y* in

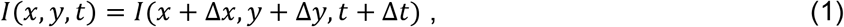

where *I*(*x*, *y*, *t*) represents the actin fluorescence intensity at frame *t*. The intensity *I* that exists at point (*x*, *y*) at time *t* translates to a new point (*x* + Δ*x*, *y* + Δ*y*) at some future time *t* + Δ*t*. Expanding about small Δ*x* and Δ*y* and neglecting second-order derivatives yields the master optical-flow equation

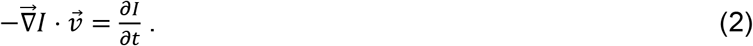

This governing equation is underdetermined, and so the Lucas-Kanade optical flow constraint^19^ was applied to calculate flow fields. This constraint prescribes that all pixels in a small window centered at (*x*, *y*) each have the same translational optical-flow vector. The equation can then be solved using the least-squares criterion (an explicit derivation is given in the **Methods Section**) to yield the intensity flow, 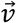(Figure 3b, center panel). Solving for 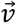 requires use of the negative spatial gradient, 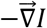 (Figure 3b, left panel), which forms a vector field oriented away from regions of highest local intensity, and the time derivative, *∂I*/*∂t* (Figure 3b, right panel) as shown in equation (2). Using the pair of images from Figure 3a as a representative example, the spatial and temporal gradients are used to calculate the optical-flow vector field, which approximates the flow of actin between the two frames, which in this example captures the translational motion on the leading edge of the cell (Figure 3c).

Optical-flow measurements of actin intensity translation enable the quantification of the pixel-scale response of actin dynamics to nanoridge topographies (Figure 4). The green-to-magenta montages of representative HL60 and MCF10A cells show dynamic and protrusive actin behavior at the leading edge of the cell (Figure 4a). Coloring the calculated flow fields based on direction relative to the nanoridges (Figure 4b) reveals the clear bias of actin wave guidance in the direction parallel to the ridges, which is consistent with the qualitative features of the montage images in Figure 4a. Measurements of the optical-flow directions on all HL60 and MCF10A cells on both flat and ridged surface topographies are shown in the histograms of Figure 4c. In both cell types, the cumulative distribution of flow in cells on flat surfaces shows no appreciable bias in any direction. However, cells on ridges exhibit a clear preference for flow along the ridge direction.

**Figure 4:**
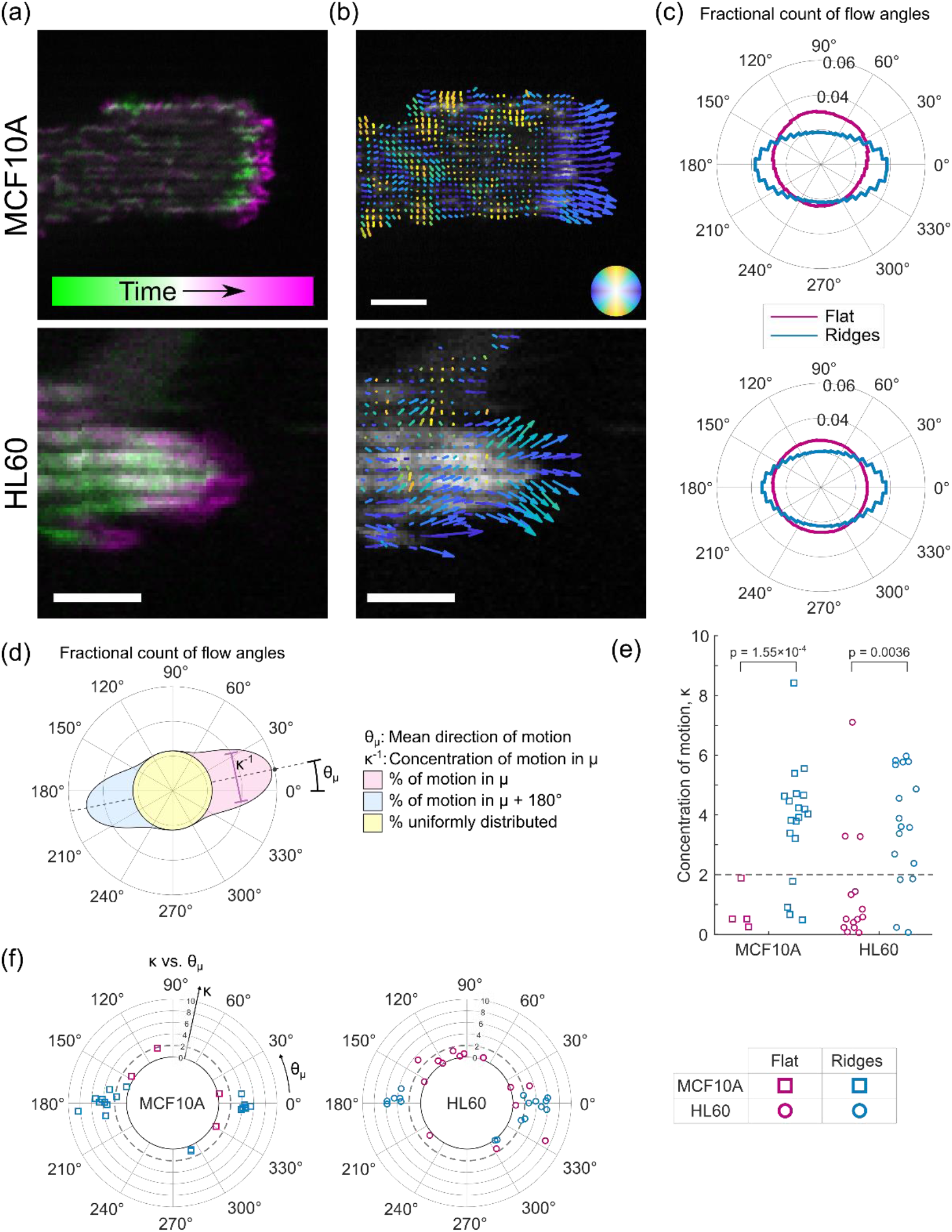
Using optical flow to measure pixel-scale guidance. (a) Representative merged time-lapse images of an MCF10A cell (top, 100 s apart) and an HL60 cell (bottom, 8 s apart). (b) Optical-flow vector fields colored by direction relative to the horizontal ridges. Blue indicates motion aligned with the ridges and yellow indicates motion perpendicular to the ridges. All scale bars are 5 µm. The dynamics in (b) are shown in **Supplemental Movies 4 and 5**. (c) Cumulative distributions of the directions of optical-flow vectors for multiple HL60 and MCF10A cells on flat and ridged surfaces; all cells are weighted equally in the distribution. [N = 4 MCF10A cells on flat surfaces from 3 independent experiments, N = 19 MCF10A cells on ridges from 4 independent experiments, N = 14 HL60 cells on flat surfaces from 2 independent experiments, and N = 17 HL60 cells on ridges from 3 independent experiments.] (d) The distribution of angles can be fit to a mixture of two von Mises distributions with five fitting parameters: *θ*_µ_ (primary direction of motion), *κ* (inversely related to distribution width), and three coefficients indicating the component of motion in the *θ*_µ_ direction, the component in the *θ*_µ_+180° direction, and the component that is uniform. (e) In both MCF10A and HL60 cells, the distribution width, parameterized by *κ*, shows significant differences (*p* = 0.000155 and *p* = 0.0036) on flat versus ridged surfaces. (f) For each cell, the mean direction of motion (angular axis) is plotted versus *κ* (radial axis). Values of *κ* less than 2 (indicated by the dashed line) indicate cells with direction distributions that are statistically indistinguishable from a uniform distribution.

For further quantification, we fit the distribution of flow directions from each cell to a bimodal von Mises model with a constant offset (**Methods Section**). The distribution used consists of a uniform component and two peaked components that are 180° apart. The five parameters of the bimodal model are illustrated in Figure 4d. The angle *θ_µ_* indicates the direction of the main component, and 1/*κ* is proportional to the width of the distribution. The values of *κ* on ridged regions are significantly higher than those than on flat regions for both the MCF10A and HL60 cells (Figure 4e, *p* = 0.000155 and *p* = 0.0036), indicating that the ridges strongly guide the actin flows in a bidirectional manner. A comparison of *κ* and *θ*_µ_ shows that cells with a bidirectional actin flow (i.e., high *κ* values) are more likely to be guided along the ridge direction (Figure 4f).

Although the optical-flow vector field indicates preferred directions of actin flow, it does not yield propagation speeds of actin polymerization waves directly. The magnitude of an optical-flow vector incorporates both the shift of actin in space and the change in actin intensity over time. This submicron-scale (i.e., pixel-scale) flux of intensity does not translate directly into characteristics of the dynamics that are notable on the micron-scale (i.e., tens of pixels), such as the organization of waves across ridges seen in neutrophils in Figure 2f or the speed of wave propagation. To quantify these micron-scale characteristics we combined similarly-oriented optical-flow vectors into clusters (Figure 5a), which were then tracked over time. To ensure that we track robust clusters, we apply additional constraints, such as requiring sufficiently large intensity changes (see **Methods Section**). The result of this clustering is the identification of broad regions of actin that move collectively (Figure 5b).

**Figure 5:**
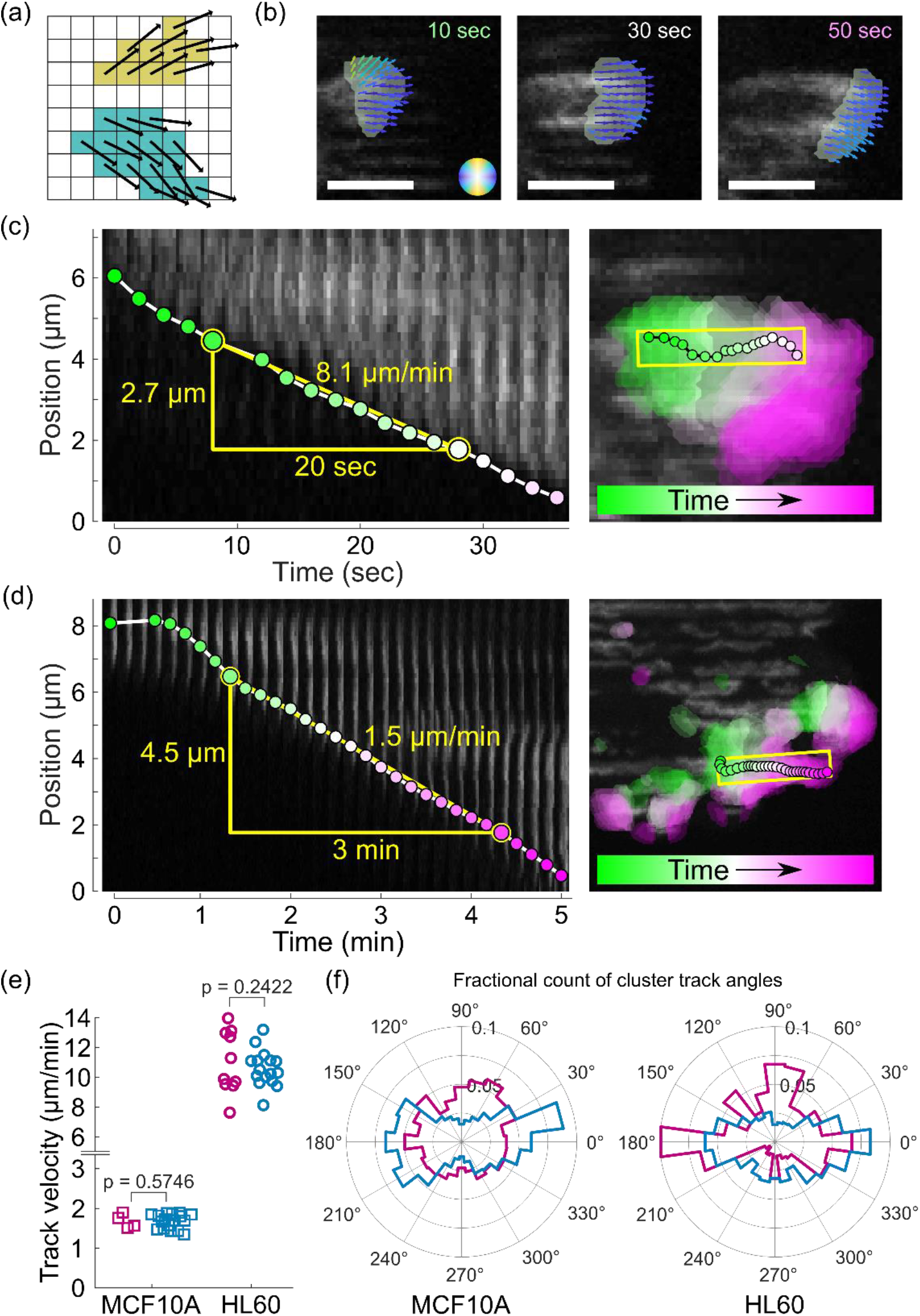
Clustering of optical-flow vectors to measure micron-scale dynamics. (a) Similar flow vectors are grouped into clusters. (b) Clusters contain optical-flow vectors with a wide array of orientations, resulting in micron-scale structures. All scale bars are 5 µm. (c, d) Particle-tracking algorithms are applied to the tracked clusters. The cluster tracks are consistent with the motion at the leading edge of the actin wave seen in the kymograph in both HL60 (c, left) and MCF10A (d, left). Panels to the right of each kymograph show the same track in the two-dimensional context of the cells; clusters found throughout the cell over time are indicated by colored regions. The dynamics in (c, d) are shown in **Supplemental Movies 6 and 7**, respectively. (e) The cluster tracks are used to determine speed distributions of actin waves on the ridges in the MCF10A and HL60 cells, which show no significant difference (*p* = 0.57 and *p* = 0.24) between flat surfaces and ridges. (f) Cumulative angle distribution of cluster track directions for multiple HL60 and MCF10A cells on flat and ridged surfaces; all cells are weighted equally in the distribution. [N = 4 MCF10A cells on flat surfaces from 3 independent experiments, N = 17 MCF10A cells on ridges from 4 independent experiments, N = 10 HL60 cells on flat surfaces from 2 independent experiments, and N = 16 HL60 cells on ridges from 3 independent experiments.]

We applied peak-finding and tracking algorithms^20^ to follow the locations of maximum alignment of these optical-flow clusters on the micron scale. Although the optical-flow results shown in Figure 4 follow motion on the pixel scale, by following the peak alignment of the optical flow, we are able to track larger coordinated clusters of actin. Our tracking is consistent with the actin dynamics seen in kymographs, as shown by representative kymographs of HL60 (Figure 5c) and MCF10A (Figure 5d) cells overlaid with the tracked location of actin waves. Unlike kymographs, which are sensitive to motion along a chosen direction, tracks of clustered flow vectors reveal the micron-scale motion of actin in two dimensions. The benefits of our approach are illustrated in Figure 5d for an MCF10A cell. For the initial frames, the kymograph indicates a stationary actin structure (Figure 5d, left), but when the actin dynamics are viewed in two dimensions (Figure 5d, right) it is evident that in the early frames this wave is moving perpendicular to the ridges, a motion that cannot be captured in this one-dimensional kymograph. Thus, the combination of optical flow, clustering, and tracking allows us to follow actin waves without being limited to tracking only motion that occurs along a straight line.

The speeds of the tracked actin clusters (Figure 5e) are similar to speeds derived from actin kymographs (Figure 5c,d, and ***Supplemental Figure 1***), despite an approximately order-of-magnitude difference in speed between the two cell types that is consistent with their distinct *in vivo* functions and with previously reported cell-migration speeds^21,22^. For both cell types we find no significant difference (*p* = 0.57 and *p* = 0.24) between actin-wave speeds on flat and ridged surfaces (Figure 5d), implying that topography steers actin dynamics but does not alter wave speeds. On flat surfaces, the directions of the clusters are distributed uniformly for MCF10A cells but show distinct peaks in multiple directions for HL60 cells. This observation is consistent with the polarized character of actin in several of the HL60 cells on flat surfaces, corresponding to *κ* values greater than 2 in Figures 4e and 4f.

## DISCUSSION

Extracellular texture, which is an important component of the three-dimensional, *in vivo* environment, is capable of spatially patterning actin and modulating actin dynamics. Using nanofabricated ridge structures in conjunction with optical-flow approaches, we are able to probe and quantify this intracellular response to extracellular textures in a systematic manner.

Previous studies of *Dictyostelium discodium*^11,12^, B cells^9^, and tumor-associated fibroblasts^23^ showed similarity in actin response to texture, which suggests that guidance of actin driven by texture (esotaxis) is broadly conserved across cell types. Controlled textures are thus a useful model microenvironment for the systematic, reproducible, and quantitative study of intracellular dynamics. Here we demonstrated the analysis of time-lapse images of two cell types that have distinct physiological function. Neutrophil-like HL60 cells are polarized and highly motile, and respond to a variety of cues as they fulfill their role in the immune system. Epithelial MCF10A cells, however, have a non-motile physiological function. Nevertheless, both cell types show clear, and quantitatively similar, actin dynamics in response to surface textures. Consistent with our prior results^11,23^, we find that nanoridges lead to persistent streaks of actin that are not seen on flat surfaces.

Optical flow enables the quantification of both the reproducible streaks of actin seen on nanoridged surfaces and the more chaotic actin waves seen on flat surfaces. The latter waves are typically much wider than guided actin waves. On flat surfaces the waves often change direction, and can also grow wider and split. Such motion phenotypes are not easily captured with standard techniques such as kymographs. Optical flow enables us to follow these dynamics, and thus yields insights beyond those derived from kymograph-based techniques. We note that optical flow is suitable for comparisons of systems imaged under different conditions (e.g., 60× vs 100× imaging and different frame rates, see **Methods Section**), enabling the comparisons of widely varying cell sizes and migration speeds.

Using submicron-scale optical flow and associated micron-scale analysis, we have shown that both MCF10A and HL60 cells have actin flows that are biased along nanoridges. By clustering similarly-oriented optical-flow vectors, we are able to measure the speed of actin waves within the cell. The measured speeds are comparable to speeds calculated from actin kymographs. Optical-flow analysis allows us to determine that the speeds do not differ significantly on flat versus ridged regions. This finding indicates that nanotopography guides but does not fundamentally alter the speed of actin dynamics. We measure actin-wave speeds on the order of 1 µm/min in the MCF10A cells, consistent with previously reported cell migration speeds of approximately 0.5 µm/min^21^. In the HL60 cells we find actin speeds ranging from approximately 8 to 14 µm/min, consistent with the 8 µm/min speed for cell migration previously reported^22^.

Fitting the optical-flow vectors to a bimodal von Mises distribution enables quantification of the differences in the directionality of actin flows on flat and ridged surfaces in both cell lines. The fit parameters also show differences in actin polarization in these two cell lines. HL60 cells occasionally exhibit coordinated and directed actin flows even on flat surfaces, whereas MCF10A cells on flat surfaces show uniform direction distributions of actin waves. On the micron scale, actin-wave tracks from individual HL60 cells on flat surfaces generally polarize and have a preferred direction, consistent with the behavior of immune cells, which tend to polarize and migrate in a directed manner. Tracks from MCF10A cells on a flat surface, on the other hand, are more directionally uniform for each cell. In both cell types, ridges guide actin waves in a bidirectional manner.

The quantitative actin responses in MCF10A and HL60 cells support a model in which surface texture provides a symmetry-breaking cue that leads to nucleation of actin polymerization. Under flat tissue-culture conditions, which lack symmetry-breaking cues, actin polymerization relies on spontaneous nucleation or edge effects^24^. Edge effects may lead to morphological features such as the lamellipodia seen in HL60 cells on flat surfaces in Figure 2b. By changing the landscape on which nucleation occurs, surface texture can lead to actin polymerization in other locations of the cell, such as the persistent streaks seen in the center of MCF10A cells on ridges in Figure 2a.

There are multiple mechanisms by which cells may respond to local forces and geometry^25^, including sensing mechanisms that can respond to membrane curvature on a variety scales^26^. In some cases, sensing mechanisms may rely on the preferential formation of focal adhesions. This hypothesis is consistent with previous results on focal adhesion localization and orientation in response to surface texture^13,14^. Although MCF10A cells form strong focal adhesions that may align with texture cues^13^, the HL60 cells form weaker adhesions, and the previously studied *Dictyostelium discodium* cells^11,12^ are not known to form integrin-mediated focal adhesions. Thus, the dominant mechanism of surface texture response likely depends on both the cell type and the extracellular environment. Other surface sensing mechanisms include cytoskeletal components such as septins, which respond to micron-scale curvatures^27^, and BAR domains, which sense nano-scale curvature^28^. Proteins with BAR domains have been linked to actin assembly^29^ as well as to key components of actin-regulating pathways such as WAVE and Rac^30,31^. Additionally, evidence suggests that topography is capable of shifting multiple gene-expression pathways^32^.

Although the systematic modulation and interrogation of all possible molecular factors of esotaxis is beyond the scope of this article, our analysis yields two remarkable constraints on the molecular sources of esotaxis. First, the speed of actin waves is not altered by esotaxis. Second, the directional guidance provided by nanotopography is comparable in the two cell types investigated, despite their disparate functions and migratory phenotypes. Quantitative analysis of esotaxis as physical phenotype could yield crucial prognostic disease insights, especially in the case of cancer, in which changes in the texture of the microenvironment correlate with disease progression.

## METHODS

### Cell Culture and Imaging

HL60 YFP-actin cells were a gift from Dr. Orion Weiner of the University of California San Francisco. The cells were cultured in RPMI 1640 medium, Glutamax (Life Technologies) supplemented with 10% heat-inactivated fetal bovine serum (Gemini Bio). Cells were passaged every 2-3 days and kept between 3×10^5^ to 1×10^6^ cells/mL. For differentiation, cell media was additionally supplemented with 1.3% dimethyl sulfoxide Hybri-Max (Sigma Aldrich) for 5 days before imaging. Actin dynamics of HL-60 YFP-actin cells were observed by confocal fluorescence and bright-field time-lapse imaging using a PerkinElmer spinning-disk confocal microscope with a water immersion 60× objective (0.21 µm/pixel). Images were recorded every 2 s. We note that this method of plating resulted in the imaging of some multicellular clusters of HL60 cells; these clusters were removed from further analysis.

Preparation for imaging included a 10 µg/mL coating of fibronectin on the substrates. Cells were plated and allowed to settle. After approximately 30 min, N-Formyl-Met-Leu-Phe (fMLF, Sigma Aldrich) was added to 1μM. HL-60 cells were imaged on flat resin and ridged nanotopographies beginning between 10-15 s after fMLF stimulation. All images analyzed in this work were obtained after fMLF stimulation.

MCF10A LifeAct-EGFP cells were a gift from Dr. Carole A. Parent. These cells were cultured in DMEM/F12 media supplemented with 5% horse serum, 10 μg/mL insulin (ThermoFisher Scientific), 10 ng/ml EGF (Peprotech, Rocky Hill, NJ), 0.5 μg/mL hydrocortisone, and 100 ng/mL cholera toxin (both Sigma, St Louis, MO). The media was additionally supplemented with 2 μg/mL puromycin dihydrochloride (ThermoFisher Scientific) to select for EGFP-positive cells. Before imaging, cells were plated on a nanoridged surface coated with collagen IV and were allowed to adhere to the surface overnight. Actin dynamics were studied by confocal fluorescence and bright-field, time-lapse imaging using a PerkinElmer spinning-disk confocal microscope with a 100× objective (0.14 µm/pixel). Images were recorded every 10 s.

For both cell types, data were collected using PerkinElmer’s Volocity software (version 6.4.0). The spinning-disk confocal microscope was equipped with a Hamamatsu ImagEM X2 EM-CCD camera (C9100-23B) which recorded 12-bit images.

### Surface Fabrication

The nanotopographies were designed and fabricated using multiphoton absorption polymerization (MAP), as described previously^13^. A drop of prepolymer resin (1:1 w/w tris (2-hydroxy ethyl) isocyanurate triacrylate (SR368):ethoxylated (6) trimethylolpropane triacrylate (SR499) (both from Sartomer), 3% Lucirin TPO-L (BASF)) was sandwiched between a coverslip and a plasma-treated microscope slide that had been functionalized with acrylate groups^13,33,34^. The coverslip was mounted onto the stage of an inverted microscope (Zeiss Axiovert 135). A beam of 150 fs pulses centered at 800 nm from a Ti:sapphire oscillator (Coherent Mira 900) was directed into the microscope and through a high-numerical-aperture objective (Zeiss alpha-Plan Fluar 100x; NA 1.45). The stage motion and shutter were controlled using a program written in LabVIEW (National Instruments). Once the pattern was fabricated, the sample was developed in ethanol and baked at 110 °C for 1 h.

A replica molding approach was then used to mold the chemically-functionalized pattern^13^. This step included the initial casting of a *hard*-poly(dimethylsiloxane) (*h*-PDMS) layer (1,000 rpm, 40 s) to better resolve nanoscale features of the topographical pattern. This layer was allowed to sit on the pattern at room temperature for 2 h, and was then baked at 60 °C for an additional 1 h. Finally, Sylgard 184 was mixed (10:1 elastomer base:curing agent) and poured onto the initial *h*-PDMS layer. The PDMS was cured at 60 °C for 1 h 10 min.

The MAP-fabricated structure was then reproduced through a soft-lithographic technique. A drop of the aforementioned resin was sandwiched between the PDMS mold and an acrylate functionalized coverslip and was then exposed to ultraviolet (UV) radiation from a lamp for a desired amount of time. After the resin cured, the coverslip was peeled off the mold. This process was repeated many times to produce enough replicas to perform the necessary experiments. The replicas were soaked in ethanol for at least 4 h and subsequently baked/dried in an oven at 110 °C for 1 h. Samples used to study MCF10A cells were also soaked in UltraPure water (ThermoFisher) for approximately 12 h.

### Kymographs

Kymographs were created in MATLAB by manually selecting a rectangular region in an actin image. Fluorescence intensities inside the region were averaged across the short axis of the region; this process was repeated for each image in the time-lapse sequence, and the resulting intensity data were combined into the kymographs shown in Figure 2 and **Supplemental Figure 1**.

### Optical Flow

The Lucas-Kanade optical-flow method^19^ was used to capture the direction and strength of intensity flow of fluorescent actin and to produce vector fields indicating actin motion. The optical-flow vector field of an image series is the field of apparent translation in the image plane, as is shown schematically in Figure 3B. Calculating the optical flow for two adjacent two-dimensional images in an image series requires solving for the unknowns Δ*x* and Δ*y* in Eq (1), as described above. The Lucas-Kanade method uses a least-squares regression approach to solve for the best optical-flow vector on a pixel-by-pixel basis under the assumption that all pixels within a “neighborhood” move in a similar direction^19^. If solving for the optical-flow vector of some point *p* with coordinate (*x*_*p*_, *y*_*p*_, *τ*), the master optical flow equation requires that the optical flow vector of point *p* and all points in the neighborhood about *p* (points 1, 2, …, *p*, …, *N*) follow the underdetermined relation

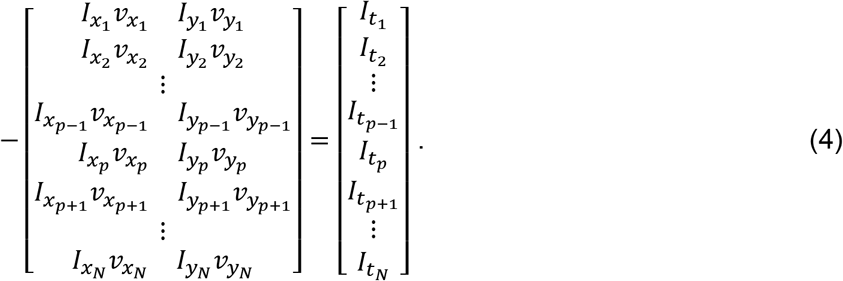

Under the Lucas-Kanade assumption, the vector for point *p* is assigned to all points in the neighborhood

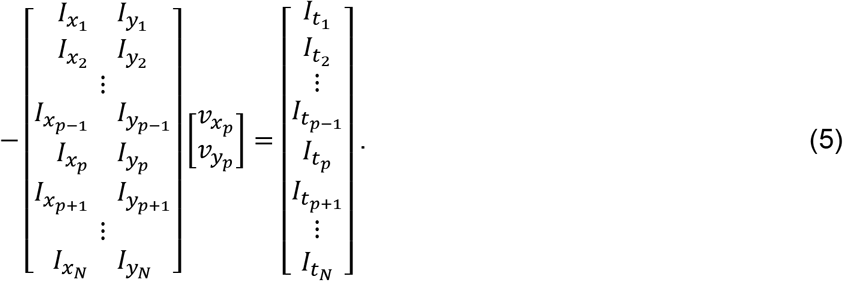

The least-squares solution to equations of this form, 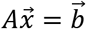, is 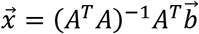. Furthermore, we implement a scheme using a Gaussian weight matrix centered at point *p* to ensure that pixels near *p* have more influence over the result. The equation then becomes

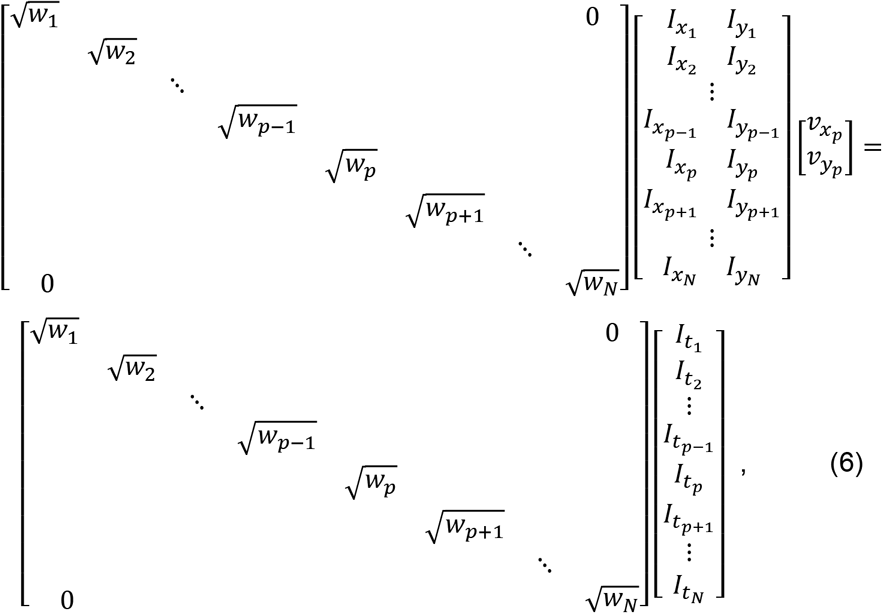

where *w* is a 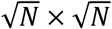 centered Gaussian matrix. The equation then takes the form 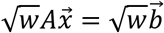 and has solution 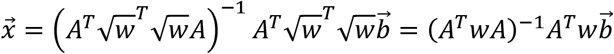.

Optical-flow reliability was defined as the smallest eigenvalue of the *A*^*T*^*wA* matrix^35,36^, and was used as a mask to remove flow vectors that were created by noise or by an ill-defined least-squares calculation. The optical-flow weight matrix for MCF10A cells was a 27 × 27 pixel Gaussian with a standard deviation of 4.5 pixels (0.63 µm). The optical-flow weight matrix for HL60 cells was 19 × 19 pixel Gaussian with standard deviation of 3 pixels (0.63 µm).

### von Mises Model of Flow Distribution

Optical-flow distributions were modelled with a modified bimodal von Mises distribution (von Mises distributions are a continuous and differentiable analogue of normal distributions on a circle with similar statistical properties). The model was defined as

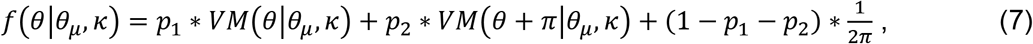

where *VM*(*θ*|*θ*_*μ*_, κ) is the von Mises distribution

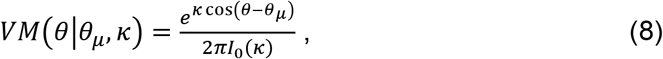

and *I*_0_(κ) is the modified Bessel function of the first kind. The maximum likelihood estimates of the parameter *κ* were used for statistical analyses.

### Cluster-Tracking Analysis

Regions of actin fluorescence were clustered using the direction of optical-flow vectors together with an optical-flow reliability threshold, and by requiring that actin intensity change over time (see ***Supplemental Figure 2*** for a visualization of this workflow). The dot products between optical-flow vectors around a point *p* (i.e. vectors *ν*_1_, *ν*_2_, …, *ν*_*p*−1_, *ν*_*p*+1_, …, *ν*_*N*_) were calculated and accumulated using a Gaussian weighting scheme to a single scalar alignment metric. The alignment metric is defined as

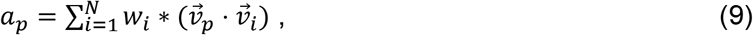

where *w* is a renormalized 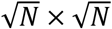 centered Gaussian matrix with a center manually set to 0. This calculation was carried out for each pixel.

To require that the actin intensity change over time a mask of the thresholded difference image between subsequent frames was calculated. For every pair of adjacent frames, *I*_*t*_ and *I*_*t*+Δ*t*_, the resulting mask took value 1 where *I*_*t*+Δ*t*_ > *I*_*t*_ and 0 otherwise. For our analysis, Δ*t* = 30 s for MCF10A and 6 s for HL60.

To calculate the final clustered regions, the alignment metric *a*_*p*_, optical-flow reliability λ_*p*_, and difference-image mask were multiplied in an element-wise fashion to create a final cluster image. The cluster image was inputted into a peak-finding algorithm to locate peaks in the resulting intensity profile, and the Crocker-Grier particle-tracking algorithm^20,37^ was used to track coordinates of the resulting peaks over time.

The clustering weight matrix for MCF10A was a 27 × 27 pixel Gaussian with a standard deviation of 4.5 pixels (0.63 µm). The clustering weight matrix for HL60 cells was 19 × 19 pixel Gaussian with standard deviation of 3 pixels (0.63 µm). The diameter of the peaks used in `pkfnd.m`^20^ was 15 pixels (2.1 µm) for MCF10A cells and 10 pixels (2.1 µm) for HL60 cells. The maximum displacement used in `track.m`^20^ was 11.5 pixels (1.54 µm) for MCF10A cells and 7 pixels (1.47 µm) for HL60 cells. Tracks measured in the movies of MCF10A cells were only considered if they were tracked for more than 3 frames (30 s) and tracks measured in movies of HL60 cells were only considered if they were tracked for more than 3 frames (6 s).

### Statistical Methods

Measurements of κ (Figure 4e) and actin-wave speed (Figure 5e) were compared on flat versus nanoridged surfaces using two-sample t-tests with unequal variances. A two-tailed t-distribution was used to calculate the reported p-values. A full description of the statistical parameters involved in these tests is provided in **Supplemental Dataset 1**.

## ACKNOWLEDGEMENTS

We thank the University of Maryland Imaging Incubator Core Facility for use of their systems in collecting images for this work. We appreciate Ema Smith’s work on the optical flow code. This work was supported by AFOSR grant number FA9550-16-1-0052. R.M.L was supported by NCI/NIH Award Number T32CA154274. L.C. was supported by COMBINE NRT Award Number 1632976 and NIH 1U01GM109887.

## AUTHOR CONTRIBUTIONS

R.M.L., L.C., J.T.F., and W.L. interpreted and analyzed the data. L.C. developed the optical flow based analysis code. M.J.H. fabricated the nanoridge surfaces. A.O. and P.A. collected the time-lapse images. J.T.F. and W.L. supervised the project. R.M.L., L.C., J.T.F., and W.L. wrote the manuscript. All authors edited the manuscript.

