## Supplemental Information for "Quantifying topography-guided actin dynamics across scales using optical flow"

#### SUPPLEMENTAL MOVIES

**Supplemental Movie 1**

Time-lapse images of the MCF10A LifeAct-EGFP cells shown in **Figure 1**. Images were taken every 10 s and are shown at 20 frames/s; the clock in the upper left corner indicates min:s. The scale bar indicates 10 µm.

**Supplemental Movie 2**

Time-lapse images of the MCF10A LifeAct-EGFP cells shown in **Figure 2**. Images on the left are bright field images corresponding to the actin shown on the right. Images were taken every 10 s and are shown at 20 frames/s; the clock in the upper left corner indicates min:s. The scale bar indicates 10 µm.

**Supplemental Movie 3**

Time-lapse images of the HL60 Actin-YFP cells shown in **Figure 2**. Images on the left are bright field images corresponding to the actin shown on the right. Images were taken every 2 s and are shown at 20 frames/s; the clock in the upper left corner indicates min:s. The scale bar indicates 10 µm.

**Supplemental Movie 4**

Time-lapse images of the optical flow results for the example MCF10A cell shown in **Figure 4b**. Actin fluorescence images are overlaid with color-coded optical-flow vectors. Blue vectors are pointing left and right, and yellow vectors are pointing up and down. Images were taken every 10 s and are shown at 12 frames/s; the clock in the upper left corner indicates min:s.

**Supplemental Movie 5**

Time-lapse images of the optical-flow results for the example HL60 cell shown in **Figure 4b**. Actin fluorescence images are overlaid with color-coded optical-flow vectors. Blue vectors are pointing left and right, and yellow vectors are pointing up and down. Images were taken every 10 s and are shown at 12 frames/s; the clock in the upper left corner indicates min:s.

**Supplemental Movie 6**

Time-lapse images of the HL60 actin track shown in **Figure 5c**. Actin images are overlaid with clusters regions and cluster tracks. The clusters are color-coded to match the times indicated in **Figure 5**. Images were taken every 10 s and are shown at 12 frames/s; the clock in the upper left corner indicates min:s. The scale bar indicates 5 µm.

**Supplemental Movie 7**

Time-lapse images of the MCF10A actin track shown in **Figure 5d**. Actin images are overlaid with clusters regions and cluster tracks. The clusters are color-coded to match the times indicated in **Figure 5**. Images were taken every 10 s and are shown at 12 frames/s; the clock in the upper left corner indicates min:s. The scale bar indicates 5 µm.

### SUPPLEMENTAL DATA

**Supplemental Dataset 1**

This file provides the statistical parameters for the t-tests conducted in this manuscript, including sample means, standard deviations, and p-values.

**SUPPLEMENTAL FIGURES**

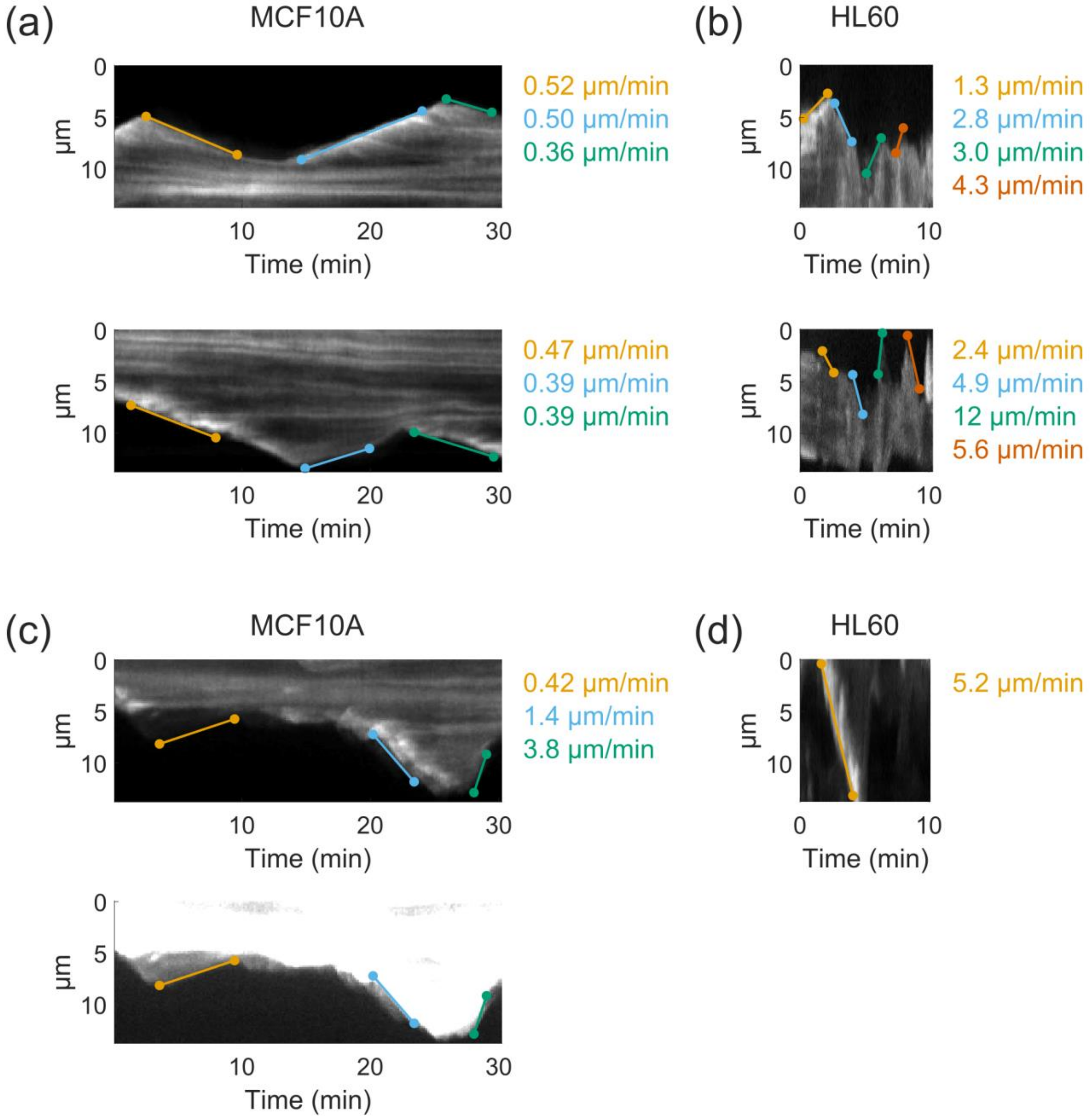


**Supplemental Figure 1: Actin-Wave Speeds**. Actin-wave speeds were calculated from the kymographs shown in ***Figure 2*** by manually selecting two points on the kymograph (using the `ginput` function in MATLAB) and calculating the slope between the points. The speeds shown here agree with the distribution of speeds found using optical-flow-based tracking (***Figure 5***) for a MCF10A on a flat surface (a), an HL60 cell on a flat surface (b), a MCF10A cell on a ridged surface (c), and an HL60 cell on a ridged surface (d). The bottom panel of (c) shows a saturated version of the underlying kymograph to emphasize the protrusions being tracked.

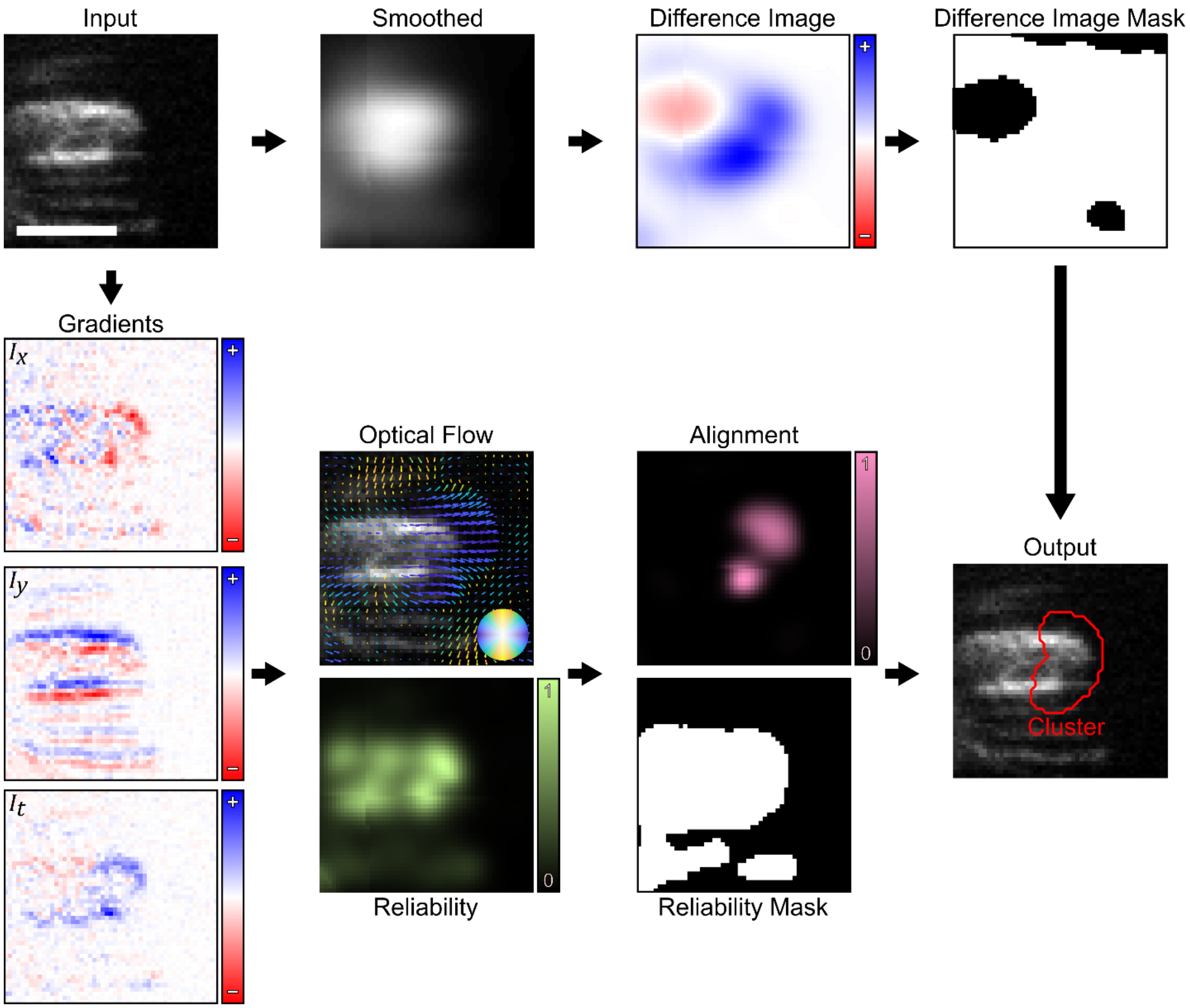


**Supplemental Figure 2: Cluster Finding Workflow**. Input time-lapse images (upper left) are used to generate a series of masks along two independent workflows:

1) To generate the difference image mask, images are smoothed and then subtracted from adjacent frames. The difference image mask is calculated by applying a threshold (in this work, the threshold is set to 0).
2) To generate the alignment and reliability mask, spatiotemporal intensity gradients are calculated and used to calculate optical flow and reliability. Alignment is calculated by taking the dot product between vectors and their local neighborhood (in this work, the neighborhood is a Gaussian with a standard deviation of 0.63 µm). The reliability mask is calculated by applying a threshold (in this work, the threshold is set to 10× the median of the reliability distribution.

Taken together, the difference image mask, alignment, and reliability mask are multiplied in an element-wise fashion to generate the final cluster image. In this work, the cluster image is used as an input to peak finding and tracking algorithms. Scale bar is 5 µm.
